# Chloroplast-localized translation for protein targeting in *Chlamydomonas reinhardtii*

**DOI:** 10.1101/2021.12.27.474283

**Authors:** Yi Sun, Shiva Bakhtiari, Melissa Valente-Paterno, Yanxia Wu, Christopher Law, Daniel Dai, James Dhaliwal, Khanh Huy Bui, William Zerges

**Affiliations:** Department of Biology, Concordia University, 7141 Sherbrooke W, Montreal, Quebec, H4B 1R6, Canada; Dept. of Plant Sciences, University of Oxford, South Parks Road, Oxford OX1 3RB, United Kingdom; Centre for Microscopy and Cell Imaging, Concordia University, 7141 Sherbrooke W., Montreal, Quebec, H4B 1R6, Canada; Department of Anatomy and Cell Biology, McGill University, 3640 University, Montreal, Quebec, H3A0C7, Canada

## Abstract

Translation is localized within cells to target proteins to their proper locations. We asked whether translation occurs on the chloroplast surface in *Chlamydomonas* and, if so, whether it is involved in co-translational protein targeting, aligned spatially with localized translation by the bacterial-type ribosomes within this organelle, or both. Our results reveal a domain of the chloroplast envelope which is bound by translating ribosomes. Purified chloroplasts retained ribosomes and mRNAs encoding two chloroplast proteins specifically on this “translation domain”, but not a mRNA encoding a cytoplasmic protein. Ribosomes clusters were seen on this domain by electron tomography. Activity of the chloroplast-bound ribosomes is supported by results of the ribopuromycylation and puromycin-release assays. Co-translational chloroplast protein import is supported by nascent polypeptide dependency of the ribosome-chloroplast associations. This cytoplasmic translation domain aligns localized translation by organellar bacterial-type ribosomes in the chloroplast. This juxtaposition the dual translation systems facilitates the targeting and assembly of the polypeptide products.

**One-Sentence Summary:** Translation is localized to a domain of the chloroplast envelope for co-translational protein targeting in *Chlamydomonas*.

## Introduction

Translation is localized in cells to ensure that the protein products get to the proper compartment, are integrated into membranes or assembled into complexes (*1*). Cytoplasmic ribosomes (cyto-ribosomes) on the ER synthesize polypeptides undergoing either co-translational import or insertion into the ER membrane. Mitochondria in yeast and human cells are bound by cyto-ribosomes which synthesize mitochondrial proteins, of which many undergo co-translational import (*2*, *3*). Mitochondria and chloroplasts contain bacteria-type ribosomes, “mito-ribosomes” and “chloro-ribosomes”, respectively, for the synthesis of proteins encoded by the small genomes in these semiautonomous organelles (*4*). Chloroplast proteins that are encoded by nuclear genes are widely believed to be synthesized at random cytoplasmic locations and undergo posttranslational import (*4*, *5*). This is based on the ability of purified chloroplasts to import *in vitro* synthesized chloroplast pre-proteins (i.e. still having their N-terminal localization sequence) and EM images of chloroplasts lacking the arrays of bound cyto-ribosomes seen on the rough ER and mitochondria (*4*). Some of the bacterial-type ribosomes within chloroplasts translate on thylakoid membranes as their nascent polypeptides undergo co-translational import (*6*). While translation on the cytoplasmic surface of a chloroplast has not been demonstrated, this possibility was raised by images from TEM and fluorescence microscopy of the unicellular green alga *Chlamydomonas reinhardtii* showing that chloroplast is adjacent to cytoplasmic region enriched in cyto-ribosomes and the mRNA encoding a chloroplast-localized protein (*7*, *8*). .

Here, we show that cyto-ribosomes translated on a “translation domain” of the chloroplast envelope in *Chlamydomonas*. These associations are demonstrated by 1) the retention of cyto-ribosomes by chloroplasts during their purification from free cyto-ribosomes and organelles known to bind them, 2) immunofluorescence (IF) microscopy images of a marker cyto-ribosomal protein (cyL4) on the chloroplast surface and 3) high resolution electron tomography images showing ribosome clusters on the outer envelope membrane. Translational activity of these chloroplast-bound cyto-ribosomes is demonstrated by results of the ribopuromycylation (RPM) and the puromycin-release assays (*9*–*12*). A proportion of these chloroplast-bound cyto-ribosomes were tethered by their nascent polypeptides, evidence that their nascent polypeptides were undergoing co-translational import *in vivo*. Synthesis of chloroplast-localized light-harvesting proteins and the small subunit of Rubisco (RBCS1 and RBCS2) on the translation domain of the chloroplast envelope is supported by results of fluorescence *in situ* hybridization (FISH) showing that purified chloroplast retained mRNAs encoding chloroplast-localized light-harvesting complex proteins (LHCPs) and the small subunits of ribulose bis-phosphate carboxylase-oxygenase (Rubisco) RBCS1 and RBCS2, but not the mRNA encoding the cytoplasmic protein ß2-tubulin. Finally, the translation domain of the envelope is spatially aligned with domains of the envelope enriched in the protein translocons of the inner/outer membrane of the chloroplast envelope (TIC and TOC) and the translation zone (T-zone), an intraorganellar compartment where chloro-ribosomes translate subunits of photosystem I (PSI) and photosystem II (PSII) of the photosynthetic electron transport chain (*8*, *13*–*15*) (Fig. 1A). Therefore, our results reveal evidence of an elaborate spatial coordination of translation of the dual translation system for photosystem biogenesis.

**Fig. 1.**
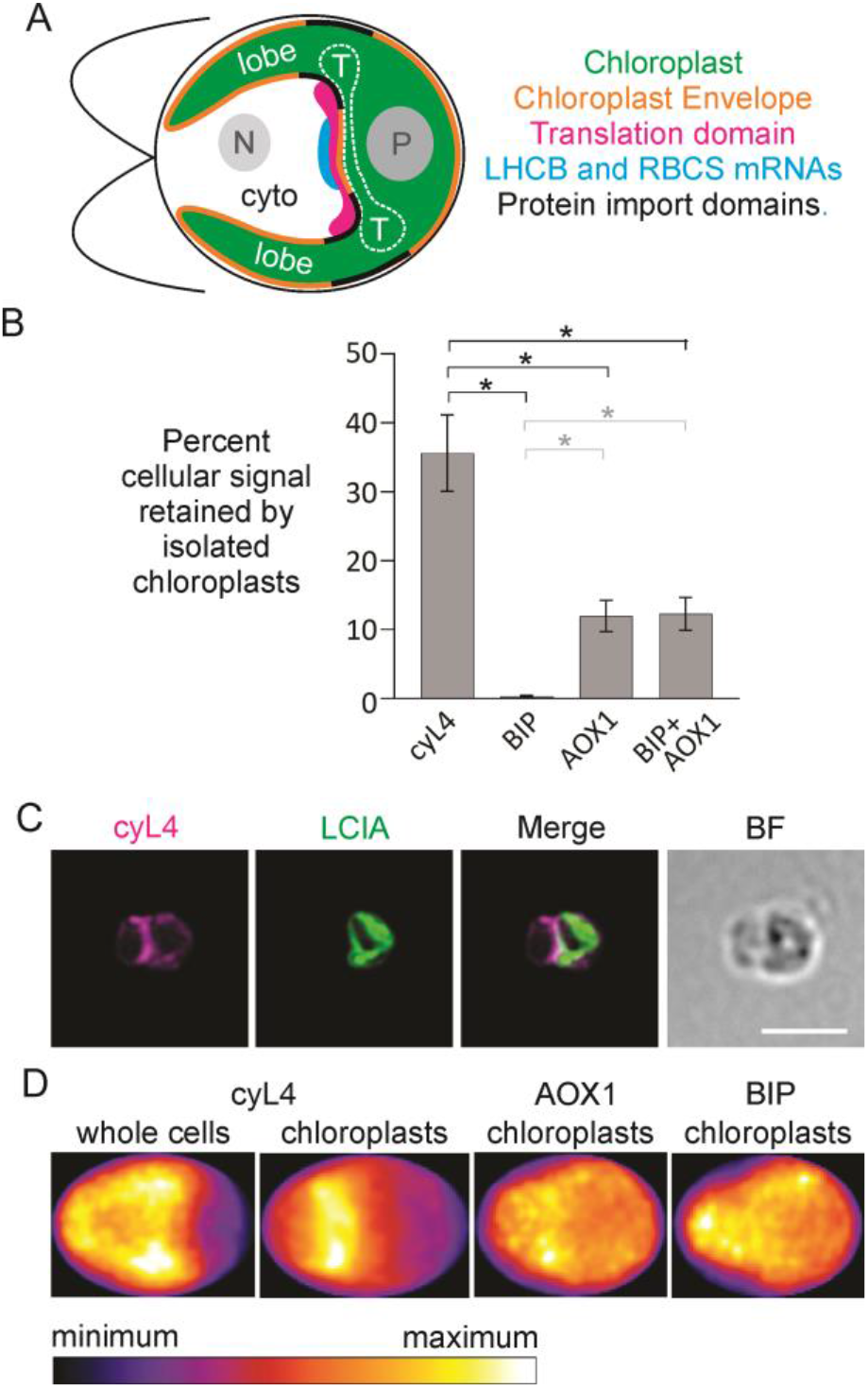
Cyto-ribosomes are bound to a domain of the chloroplast envelope. (A) An illustration shows a *Chlamydomonas* cell with the nucleus (grey sphere), cytosol (cyto) and chloroplast (green). The chloroplast has lobes which enclose the nucleus (grey sphere), and cytosol (cyto), pyrenoid (black sphere), the T-zone (T) and it is surrounded by a dual membrane envelope (orange). The translation domain of the envelope (magenta) is adjacent to the mRNA-enriched region (cyan) and overlaps envelope domains shown previously to be enriched in the TOC/TIC protein import translocons (black) (*14*). (B) Results of immunoblot analyses of marker proteins in extracts of whole cells versus isolated chloroplasts reveal that cyto-ribosomes (cyL4) preferentially copurify with chloroplasts (AtpB) relative to the organelles known to be bound by cyto-ribosomes; ER (BIP), mitochondria (AOX1). (Immunoblot results are in Fig. S4. Error bars= 1.0 SEM, n=3 biological replicates from independent cultures). (C) IF-microscopy images of purified chloroplasts show cyL4 localized to a domain of the envelope (LCIA). The absence of LCIA signal from the lobes of the chloroplast does not reflect a change in chloroplast morphology during isolation (Fig. S3). (BF, bright field, size bar, 5.0 μm) (D) Heat maps show average IF signals of cyL4, AOX1, and BIP from all cells or chloroplasts in representative data sets (CyL4, n= 32 chloroplasts or n= 102 cells; AOX1, n=22 chloroplasts; BIP48, n=48 chloroplasts).

## RESULTS

### Cyto-ribosomes are bound to a translation domain of the chloroplast envelope

To explore the possibility that translation is localized to the outer envelope membrane of the chloroplast in *Chlamydomonas*, we asked whether cyto-ribosomes copurify with chloroplasts during their isolation from away from cyto-ribosomes that are free or bound to contaminating ER and mitochondria. Isolated chloroplast retained more cyto-ribosomes than can likely be explained by contamination by mitochondria and ER (Fig. 1B). To determine whether cyto-ribosomes were bound to the purified chloroplasts, we imaged the ribosomal protein cyL4 by IF microscopy. On purified chloroplasts, the cyL4 IF signal was seen adjacent to the chloroplast envelope, which was co-IF-stained for the envelope marker protein LCIA (*16*). The cyL4 signal was strongest at a region of the chloroplast envelope bordering the central nuclear-cytosolic region (Fig. 1A and C). This localization pattern can be seen in a representative chloroplast and in the average signal distribution in all chloroplasts of the data set, but not in images of whole cells, where the signal was throughout the cytoplasm (Fig. 1C and D, Fig. S1A). This chloroplast-localized cyL4 IF signal was not from cyto-ribosomes bound to ER or mitochondria that were retained by these chloroplasts because marker proteins for these organelles did not show the same pattern as cyL4 (Figs 1D, S2A and B). These results support associations of cyto-ribosomes with a translation domain of the chloroplast envelope which spatially aligns with the T-zone within this organelle (Fig. 1A).

### Cyto-ribosomes on the translation domain of the chloroplast envelope were imaged by high-resolution electron tomography

The evidence cited against chloroplast-localized translation includes EM images of chloroplast envelope devoid of bound ribosomes and chloroplasts surrounded by a cyto-ribosome-free zone (*17*–*19*). Therefore, to determine whether cyto-ribosomes can be visualized on the chloroplast envelope, and to validate the cyL4 IF signal as a marker for them, we imaged cells with three-dimensional high-resolution electron tomography (Fig. 2). For reference, we imaged the envelope of chloroplast lobes, which did not strongly IF-stain for cyL4. The results show the presence of cyto-ribosome clusters on the chloroplast envelope domain where we observed the localized cyL4 IF signal (Compare Figs. 1C and 2C-F). Cyto-ribosome density was lower on other regions of the chloroplast envelope, e.g., of the chloroplast lobe (Fig. 2F and G). This illustrates that, cyto-ribosomes are on the chloroplast envelope, thereby, corroborating the results of IF microscopy.

**Fig. 2.**
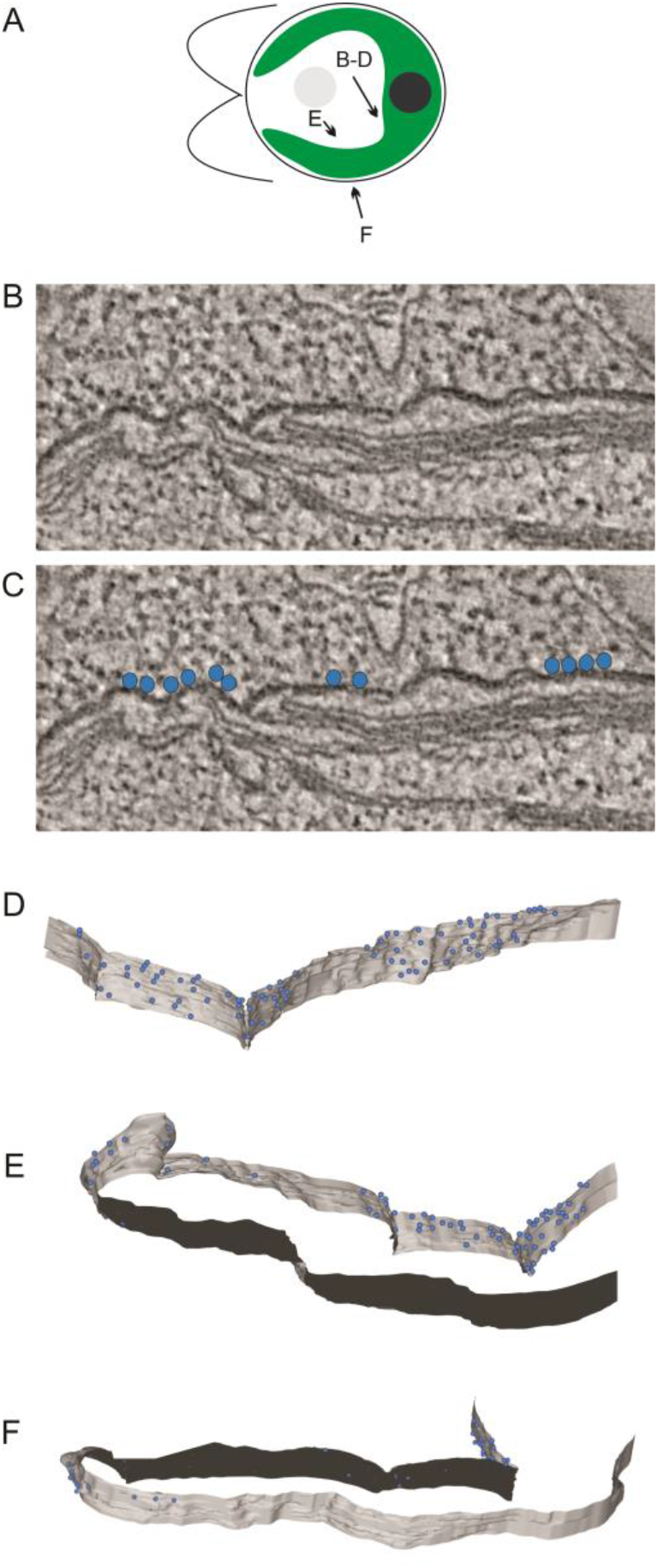
Electron tomograms show cyto-ribosomes on the outer membrane of the chloroplast envelope. (A) The illustration shows the regions where the tomographs were acquired for B-E. (B) A tomographic slice showing the region of chloroplast envelope bound by cyto-ribosomes as seen by IF microscopy (Fig. 1C). (C) The image in B with blue dots marking the cyto-ribosomes that are on the envelope. (D-E) Models of chloroplast envelope (grey, cytoplasmic face of the outer membrane; black, stromal face of the inner membrane) and bound cyto-ribosomes (blue dots) as seen from the angles shown in Panel B.

### Chloroplast-bound cyto-ribosomes are active

We used two methods to determine whether the chloroplast-bound cyto-ribosomes are translationally active. The RPM method takes advantage of the conjugation of puromycin to the nascent polypeptide when it terminates translation by imaging the IF signal from the resulting puromycin-conjugated nascent polypeptides as markers *in situ* for locations of translation *in vivo* (*12*). Routing of the puromycin-conjugated nascent polypeptides from sites of their synthesis, a concern when live cells are treated prior to fixation and IF-staining (*20*), is unlikely because we treated isolated chloroplasts with puromycin.

Chloroplasts were isolated, treated with puromycin, IF-stained with an antibody specific to puromycin, and imaged by epifluorescence microscopy. (Specificity of the puromycin signa is demonstrated in Fig S2C). Isolated chloroplasts showed the strongest IF-signal of puromycin-conjugated nascent polypeptides at the envelope domain marked by the localized cyL4 IF-signal (Fig. 3A). Localization of the puromycin signal on the cytoplasmic side of the chloroplast envelope (LCIA) demonstrates that these nascent polypeptides were from cyto-ribosomes and not chloro-ribosomes (Fig. 3B). Moreover, this puromycin-nascent polypeptide localization pattern was seen in the average signal distribution in maximal intensity projection of all chloroplasts in the data set (Fig. 3C). These results support translational activity of the chloroplast-bound cyto-ribosomes *in vivo.*

**Fig. 3.**
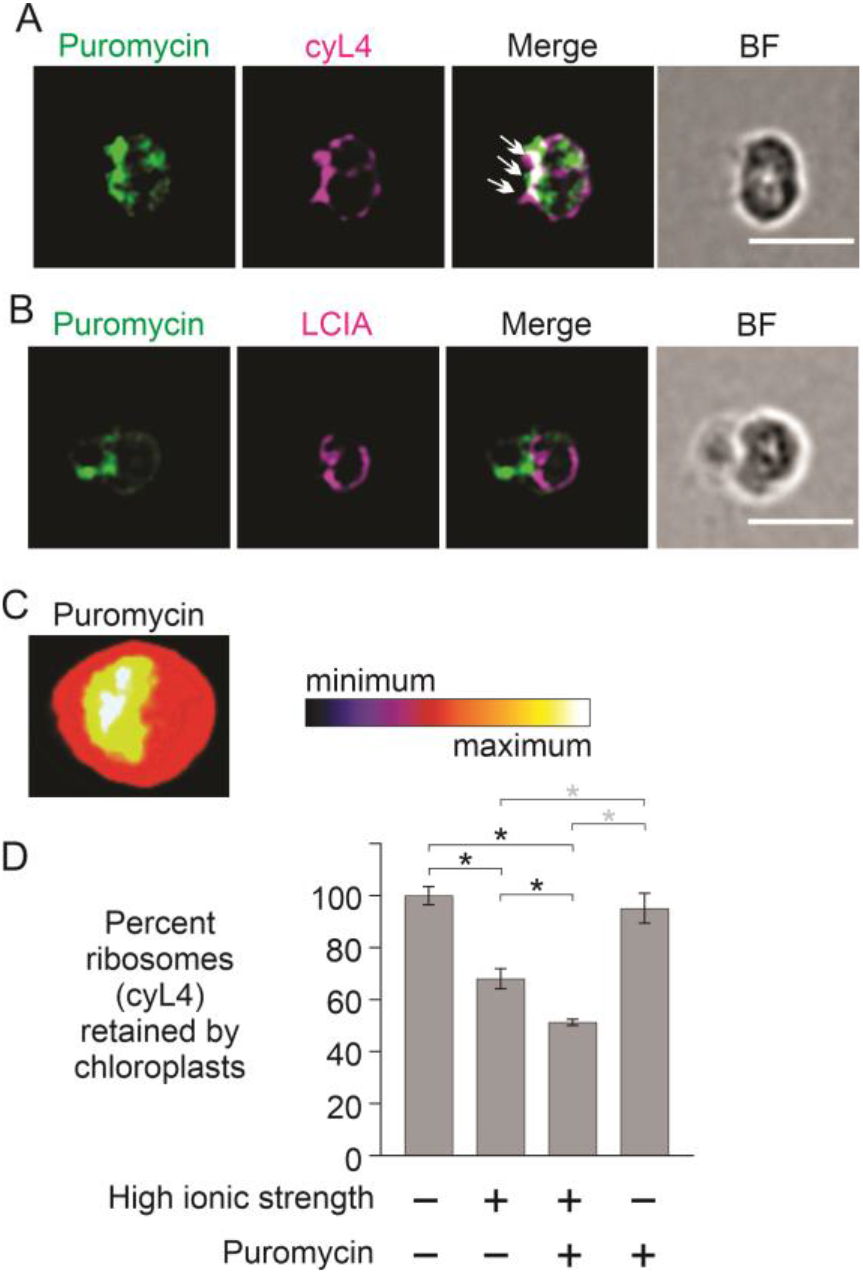
The cyto-ribosomes on the chloroplast are translationally active and tethered by nascent polypeptides. (A and B) Results of the RPM method show IF signal of the puromycin-conjugated nascent polypeptides (green), as markers of translation, localized to (A) the cyto-ribosome (cyL4) IF signal (B) on the cytoplasmic side of the chloroplast envelope (LCIA) (size bar, 5.0 μm). Arrows indicate sites of colocalization of puromycin-conjugated nascent polypeptides and cyto-ribosomes. The green IF signal is specific to puromycin (Fig S2C). (C) A heat map of the average IF signal from the puromycin-conjugated nascent polypeptides from all chloroplasts in this data set (n= 30) shows that the individual chloroplasts are representative. (D) Bar heights indicate the average proportion of cyto-ribosomes (cyL4) retained by isolated chloroplasts following the treatments indicated. (Immunoblot results represented by this graph are presented in Fig S4.) High ionic strength was 750 mM KCl. (Error bars= 1.0 SEM, n= 3 biological replicates from independent cultures).

The puromycin-release assay tests for organelle-localized translation by exploiting the specificity of puromycin for releasing translating ribosomes from their nascent polypeptides (*9*). Puromycin-induced release of cyto-ribosome from an isolated organelle is evidence that the ribosomes were translating and tethered by nascent polypeptides undergoing co-translational passage via the protein translocons in the organellar membrane (*9*, *21*). In addition, ribosomes on the ER, mitochondria and thylakoid membranes required high-ionic strength (300-750 mM KCl) to be released, because they are bound to ribosome receptors on the organelle surface (*10*, *22*, *23*). When chloroplasts were incubated in the high ionic strength condition (750 mM KCl), a significant proportion of cyL4 was released (32%, p= 0.037) (Fig. 3D). Therefore, this proportion of the cyto-ribosomes on the translation domain of the chloroplast envelope were bound by non-covalent bonds alone. Treatments with both puromycin and high ionic strength released 49% of cyL4 (p= 0.012), 17% more than were released during treatment with high ionic strength alone (p= 0.023). Therefore, these cyto-ribosomes were bound by both non-covalent bonds and their nascent polypeptides. This result confirms that some of the chloroplast-bound cyto-ribosomes were translationally active *in vivo*. It also reveals that at 17% of the ribosomes were associated by their nascent polypeptides. Similar results revealed previously that nascent polypeptides undergo co-translational import into the ER and mitochondria (*10*, *22*, *23*). Moreover, the retention of the puromycin-conjugated nascent polypeptides by the chloroplast envelope domain is consistent with their being anchored in the chloroplast envelope, for example, possibly reflecting co-translational import. (Fig. 3A-C). Finally, puromycin alone did not release a significant proportion of cyL4 (p= 0.603), revealing that few, if any, cyto-ribosomes were associated with the chloroplast by nascent polypeptides alone. That approximately 50% of the cyto-ribosomes were not dissociated by any of the treatments could reflect high affinity ribosome-chloroplast associations, ribosomes trapped within contaminating unbroken cells or both. Together, these results reveal the chloroplast bound cyto-ribosomes are active and bound to the chloroplast by both non-covalent bonds and their nascent polypeptides.

### The translation domain of the chloroplast envelope is bound by mRNAs encoding chloroplast-localized LHCB and RBCS proteins

The results above support localized translation by chloroplast-bound cyto-ribosomes for protein import into the T-zone within the chloroplast (Fig. 1A). This predicts that the translation domain of the chloroplast envelope is associated with mRNAs encoding chloroplast proteins, but not mRNAs encoding non-chloroplast proteins. We used FISH to test this prediction (*24*). The imported chloroplast proteins include subunits of the light harvesting complexes (LHCs), which each have three hydrophobic transmembrane domains and are embedded in the membranes of photosynthetic thylakoid vesicles where they associate with PSI and PSII (*25*, *26*). One might expect LHCPs to be synthesized by chloroplast-bound cyto-ribosomes and undergo co-translational import and membrane insertion because the vast majority of such hydrophobic integral membrane proteins use this targeting mechanism to prevent their misfolding and aggregation in the aqueous cytoplasm and, consequentially, impaired import and toxicity (*2*, *3*, *27*). Therefore, we asked whether chloroplasts released from cells retain mRNAs encoding LHCPs (*28*). Our FISH probe sequences are complementary to the mRNAs of *LHCBM2* (Cre12.g548400) and *LHCBM7* (Cre12.g548950), highly similar paralogues in the *LHCB* gene family (Table S1). The mRNAs detected by these probes are referred to collectively as “*LHCBM*” here. In cells, the *LHCBM* FISH signal was detected from the cytosol, where it was enriched near the chloroplast, as was reported previously (Fig. 4E and Fig S1B) (*8*). Chloroplasts retained 96% of average cellular signal, and individual chloroplasts showed localized signal closely adjacent to, but not overlapping, the chloroplast-localized cyL4 IF signal (Fig. 4A and B). Consistency of this localization pattern across all chloroplasts imaged was seen in a display of the average *LHCBM* mRNA FISH signal distribution (Fig. 4E). While the translation domain extends along the envelope between opposing lobes, the strongest average *LHCBM* mRNA FISH signal was localized at the center of this domain (contrast cyL4 in Fig. 1D versus *LHCBM* and *RBCS* in Fig. 4E, illustrated in Fig. 1A). These results reveal a physical association of *LHCBM* mRNAs with the translation domain of the chloroplast envelope. We also imaged the mRNAs of *RBCS1* and *RBCS2,* which encode the small subunits of Rubisco, a chloroplast-localized enzyme (Cre02.g120100 and Cre02.g120150). We refer to these mRNAs as “*RBCS*” because our FISH probes hybridize to both (Table S1). In cells, localization of the *RBCS* mRNAs in the cytosol was not evident in most images, as was reported previously (Fig. S1C) (*8*). However, the average *RBCS* FISH signal from all cells imaged revealed localization to the approximate location of the cyto-ribosomes on the translation domain of the chloroplast envelope (Fig. 4E). Moreover, an association of the *RBCS* mRNAs with the chloroplast was revealed by our findings that free chloroplasts retained 80% of the cellular *RBCS* FISH signal and that individual chloroplasts showed this signal localized at the middle of the translation domain (marked by cyL4), like the localization of the *LHCBM* mRNA FISH signal (Fig. 4A and C). A heatmap of the average *RBCS* FISH signal confirmed this localization pattern and revealed that more of the *RBCS* mRNA FISH signal was around the entire basal (posterior) region of the chloroplast, than was the *LHCBM* FISH signal (Fig. 4E). These results support the translation of at least a few cytoplasmic mRNAs encoding chloroplast proteins by the cyto-ribosomes in the center of the translation domain of the chloroplast envelope.

**Fig. 4.**
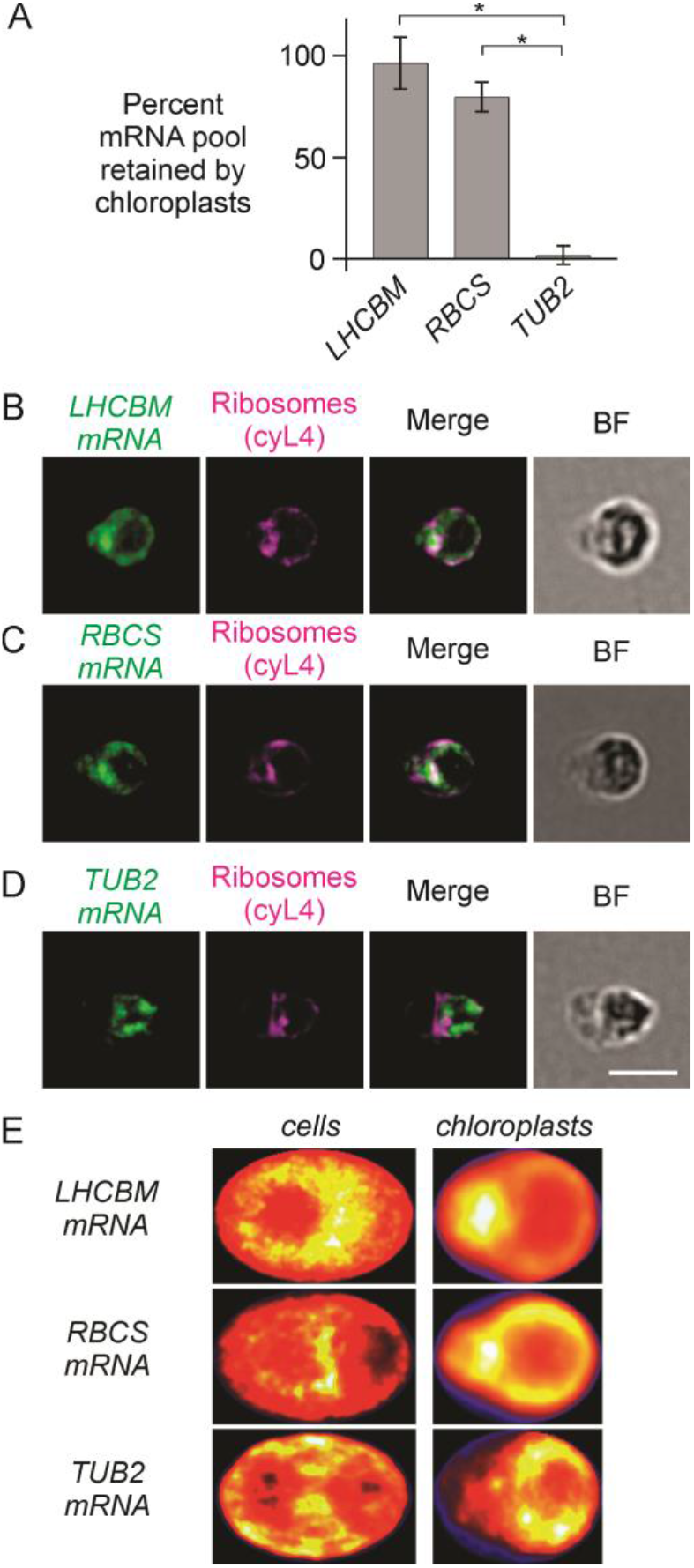
FISH results reveal that mRNAs encoding specifically chloroplast-localized proteins are bound to isolated chloroplasts. (A) Bar heights represent percentages of the average FISH signal intensities of whole cells that were retained by chloroplasts for the mRNAs indicated. (Error bars= 1.0 SEM). (B-D) Chloroplasts IF-stained for cyto-ribosomes (cyL4) and FISH-probed for the mRNAs encoding chloroplast-localized proteins of (B) the *LHCBM* mRNAs or (C) the *RBCS* mRNAs. (D) Chloroplast FISH-probed for the *TUB2* mRNA as a control mRNA encoding a non-chloroplast protein. Size bar, 5.0 μm. (E) Heat maps show the distributions of the average FISH signals in maximal intensity projections of image stacks from all cells or chloroplasts in each data set (n ≥ 30 cells or chloroplasts per data set).

To assess for specificity of chloroplast localization of mRNAs encoding chloroplast proteins, we visualized the FISH signal from the *TUB2* mRNA, which encodes ß2-tubulin, a protein of the cytoplasm and cilia (Cre12.g549550) (*29*). In cells, strong *TUB2* FISH signal was detected throughout the cytosol, as reported previously (Fig. 4E, Fig. S1D) (*7*). Chloroplasts retained 2% of this signal (Fig. 4A) which was not enriched at or near the translation domain (Fig. 4D and E). These results support chloroplast-localized translation specifically of mRNAs encoding proteins of the chloroplast.

## DISCUSSION

Our results reveal localized translation of mRNAs encoding chloroplast proteins at a domain of the chloroplast envelope in *Chlamydomonas*. This translation domain contradicts the long-standing model that all chloroplast proteins are synthesized throughout the cytoplasm (*4*). In addition, the nascent polypeptide dependency of cyto-ribosome associations with the chloroplast supports co-translational import of chloroplast proteins (Fig. 3D). In this mechanism, the emerging nascent polypeptide passes through the chloroplast envelope via the TOC/TIC translocons during its synthesis, thereby tethering the cyto-ribosome to the chloroplast. Translation localization at the ER and mitochondria, in addition to tethering by nascent polypeptides, involves cyto-ribosome receptors on the organellar surface. These receptors were revealed by requirements for high ionic strength for ribosome dissociation from these organelles *in vitro* (*23*, *30*–*32*). The possibility that cyto-ribosomes bind to receptors on the chloroplast surface is suggested by our finding that high ionic strength is required for their dissociation (Fig. 3D).

Our results reveal that chloroplast protein synthesis and import are organized spatially in a fashion analogous to mitochondrial protein synthesis in *Saccharomyces cerevisiae* and humans (*33*– *37*). Mito-ribosomes synthesize subunits of the complexes of the respiratory electron transport system and ATP synthase into the inner membrane where it invaginates to form cristae, i.e. cristae junctions (*36*–*38*). Cristae junctions are also preferential sites of the early steps of respiratory complex assembly (*36*–*38*). As such, cristae junctions are analogous to the T-zone of the chloroplast (*15*). The translation domain of the chloroplast envelope is analogous to patches of mitochondrial outer membrane located outside cristae junctions, which are bound by translating cyto-ribosomes and enriched in the mitochondrial protein import translocons (*21*, *33*, *36*, *39*). Therefore, spatial coordination of translation on and within each of the semiautonomous organelles might be a fundamental aspect of their biogenesis. This localized co-translational import of mitochondrial inner membrane proteins is hypothesized to facilitate their integration into the membrane and assembly with the locally synthesized protein products of mito-ribosomes (*32*, *40*–*42*). Similarly, we hypothesize that LHCPs, and possibly other chloroplast proteins, are synthesized at the translation domain of the chloroplast envelope and undergo co-translational import into the T-zone to facilitate their insertion into developing thylakoid membranes and their assembly with subunits synthesized by chloro-ribosomes. In this model, the homologues of the chloroplast SRP system, cpSRP43 and cpSRP54, engage the nascent polypeptide as it emerges from the TIC translocon and direct it to the translocon for co-translational insertion into developing thylakoid membranes in the T-zone, thereby obviating the proposed post-translational roles of the chloroplast SRP system.

## Acknowledgments

For technical assistance, infrastructure and support we thank the Centres for Microscopy & Cell Imaging and Structural & Functional Genomics (Concordia University) and Jeannie Mui and the Facility for Electron Microscopy Research (McGill University). For generous gifts of antibodies, we thank Prof. Hideya Fukuzawa (LCIA, Kyoto University), Dr. Elizabeth Harris (αAtpB and αcyL4, Duke University), Drs Pierre Crozet and Stephane Lemaire (αPRK) and Dr. Jonathan Yewdell for depositing PMY-2A4 at Developmental Studies Hybridoma Bank).

## Author contributions

Conceptualization: YS, SB, WZ

Methodology: YS, SB, CL, YW, MVP, KHB

Investigation: YS, SB, YW, DD, MVP, JD.

Visualization: SB, YS, CL, DD, KHB.

Funding acquisition: WZ

Project administration: WZ

Supervision: YS, KHB, WZ

Writing – original draft: WZ

Writing – review & editing: YS, SB, WZ, KHB.

## Data and materials availability

All data, code, and materials used in the analysis must be available in some form to any researcher for purposes of reproducing or extending the analysis. Include a note explaining any restrictions on materials, such as materials transfer agreements (MTAs). Note accession numbers to any data relating to the paper and deposited in a public database; include a brief description of the data set or model with the number. If all data are in the paper and supplementary materials, include the sentence “All data are available in the main text or the supplementary materials.”

## Supplementary Materials

Materials and Methods

Figs. S1 to S4

Table S1

