## Supplemental Information for "Chloroplast-localized translation for protein targeting in *Chlamydomonas reinhardtii*"

### Supplementary Materials

**This PDF file includes:**

Materials and Methods
Figs. S1 to S4
Table S1
References (*40*–*48*)

**Materials and Methods**

Strains. *Chlamydomonas reinhardtii* strain used for most experiments has a cell-wall deficiency (CW15) (CC-400 *MT*+). For the results in Figure 2, wild-type strain CC-125 was used. Strains were cultured to $1\times10^5$ cells·ml$^{-1}$ in high salt minimal (HSM) medium (*43*), with aeration, illuminated by banks of red and blue LEDs at 150 µE·m$^{-2}$·s$^{-1}$ at 23 ºC with orbital shaking (120 rpm). CC-400 was cultured in HSM containing 1.0% (w/v) sorbitol. Cultures were entrained under alternating cycles of 12 h light:12 h dark for 3 days to a density of ~$4 \times 10^6$ cells·ml$^{-1}$. Cells were harvested at the fourth hour of final light cycle by centrifugation (3,000 g, 5 min at RT). Chloroplast isolation was performed as described previously with the following modifications (*44*, *45*). Cell pellets were resuspended to $1\times10^8$ cells·ml$^{-1}$ in isolation buffer (IB) [300 mM Sorbitol, 50 mM HEPES-KOH pH 7.5, 25 mM $MgCl_2$, 0.1% (w/v) BSA]. Saponin (Sigma, # 47036) 10% (w/v) freshly dissolved in IB) was added to 0.4% (w/v), followed by incubation at 22 ºC for 10 min with occasional gentle agitation. The resuspension was passed twice through a 27-gauge needle at 0.1 mL·s$^{-1}$. Cells and chloroplasts were collected by centrifugation at 750 g for 2 min at 4 ºC. The pellet was resuspended in IB and overlaid on the Percoll gradient (*45*), which was centrifuged for 25 min at 3,200 g. The material at the 45-65% interface was collected. The Percoll was diluted by addition of 4 vol IB. Chloroplasts were pelleted by centrifugation (670 g, 1 min, 4 ºC), resuspended in buffer according to the downstream use.

Immunoblot Analysis. For the immunoblots in Fig 1E, same number of isolated chloroplasts were resuspended in SDS-PAGE loading buffer and resolved by SDS-PAGE (12% acrylamide) (*46*). Proteins were transferred to a membrane of PVDF (BIO-RAD) or, for AOX1 detection, nitrocellulose (BIO-RAD) and reacted with primary and secondary antibodies as described previously (*46*). The primary antibodies were: α-BIP (Santa Cruz sc-33757) (1:150), α-AOX1 (Agrisera AS06 152, 1:150,000), α-cyL4 (1:6,000)(*47*) and α-AtpB (1:6,000). The secondary antibody was horseradish peroxidase-conjugated goat anti-rabbit IgG antibody (KPL). Signals were detected using an ECL substrate (Thermo-Fisher) with an Imager 600 (Amersham/GE) according to the manufacturer's protocols. Signal quantification was conducted with Imager 600 Analysis Software (Amersham).

IF-staining. FISH and microscopy IF-staining was performed as described previously (*48*). The primary antibodies and the dilutions were: αcyL4 (1:1000) (*47*), αAOX1 (1:1,200), and αBIP (1:100). The secondary antibody was AlexaFluor568 conjugated to goat anti-rabbit IgG (Thermo Fisher). For dual IF-staining (Figure 1C), chloroplasts were first reacted with α-LCIA (1:700) and then indirectly IF-labelled by AffiniPure Fab Fragment Donkey Anti-Rabbit IgG (H+L) conjugated to AlexaFluor488 (Jackson ImmunoResearch Inc). Chloroplasts were reacted with αcyL4 (1:1000) and then indirectly IF-labelled by goat anti-rabbit IgG conjugated to AlexaFluor568 (Thermo Fisher). For consistency, the cyL4 IF signal is presented in magenta and other signals in green. Microscopy was carried out with a Leica DMI6000B inverted epifluorescence microscope with a 63x Plan Apo objective (NA 1.4) and further magnified by a 1.6x tube lens. Images were acquired on a Hamamatsu Orca R2 C10600-10B camera controlled by Volocity software (Improvision). Filters: Texas Red (562/40nm ex: 624/40nm em) for AlexaFluor568 and GFP (472/30nm ex: 520/35nm em) for AlexaFluor488. Acquired images were taken using Z plane stacks with a spacing of 0.2 µm per section; exposure settings, gain, and excitation intensity were kept constant where comparisons between intensities was required. For deconvolution, Z-stacks were taken by series capture at a thickness of 0.2 µm per section and were deconvoluted with AutoQuant X3 (Media Cybernetics Inc). FISH was carried out as described previously, except that of BSA concentration in hybridizations was 4.5 mg/ml (*24*). For the results in Figure 4A, the average FISH signal intensity from the probe with random sequence obtained for cells or chloroplasts in each trial was subtracted from the average intensities of the mRNA FISH probes. p-values are from 2-tailed Student's t-tests comparing, $n \geq 3$ biological replicates using independent cultures. The images in Figure 4B-D were adjusted to best show distributions of each mRNA signal using Photoshop (Adobe). The TUB2 mRNA FISH signal was seen previously throughout the cytoplasm in deflagellated cells, as we observed here, possibly because the strain that we used lacks flagella (*7*) (Supplemental Figure 1D, Figure 4D). Specificities of the FISH signals of the RBCS1/RBCS2 and LHCBM2/7 mRNAs were demonstrated previously (*8*). To determine the average distribution of fluorescent signals in cells or chloroplasts of a data set, we used a macro which operates within ImageJ, described previously (*15*, *49*) (https://github.com/Zergeslab/cellHarvester).

High resolution electron tomography. Sections of 300 nm thickness from the resin-embedded cells above were collected on Formvar support slot grids and stained. The dual-axis tilt series were collected using the FEI Tecnai $G^2$ F20 200 kV TEM equipped with a Gatan Ultrascan 4000 4k x 4k CCD Camera System Model 895 and a single tilt holder. Tilt series were then acquired at 2° increment from −60° to 60°, at 19000x magnification, 5.91 Å pixel size using SerialEM (*50*). For the second axis tilt series acquisition, the slot grid was rotated 90° manually and the same area of interest was searched manually. The dual-axis tomograms were reconstructed from the tilt series using IMOD software package (*51*). The modelling and visualization of the membrane and cyto-ribosomes were done also by IMOD.

RPM and puromycin-release assays. For the RPM method, isolated chloroplasts ($1\times10^8$ $ml^{-1}$ in IB were treated with 1.0 mM puromycin (Bioshop) for 10 min at RT and then IF-stained with a mouse monoclonal antibody against puromycin (DSHB Hybridoma Product PMY-2A4, deposited by J. Yewdell). The IF signal was specific (Fig. S2C). The puromycin-release assay followed protocols that were used to show ribosome association to ER, mitochondria and thylakoid membranes with following modifications (*10*, *22*, *23*). Cells of CC-400 ($9 \times 10^8$) were pelleted by centrifugation at 3,000 x g, 5 min at RT. Cell density was adjusted with HSM+1% sorbitol to $1.2 \times 10^7$. Chloroplasts were resuspended with 1.0 mL IB (150 µL), pelleted at 1,000 x g for 3 min at RT, resuspended with 1.0 mL of one of the following four conditions: 1) IB+ 5 mM DTT, 2) IB+ 5 mM DTT+ 750 mM KCl, 3) IB+ 5 mM DTT+ 1 mM puromycin + 750 mM KCl and 4) IB+ 5 mM DTT+ 1 mM puromycin. Samples were incubated at RT for 20 min. Chloroplast were pelleted by centrifugation (1,000 x g for 3 min at RT) for immune-blot analyses. Results are from three concurrent biological replicate experiments (i.e. from independent cultures, Fig. S4).

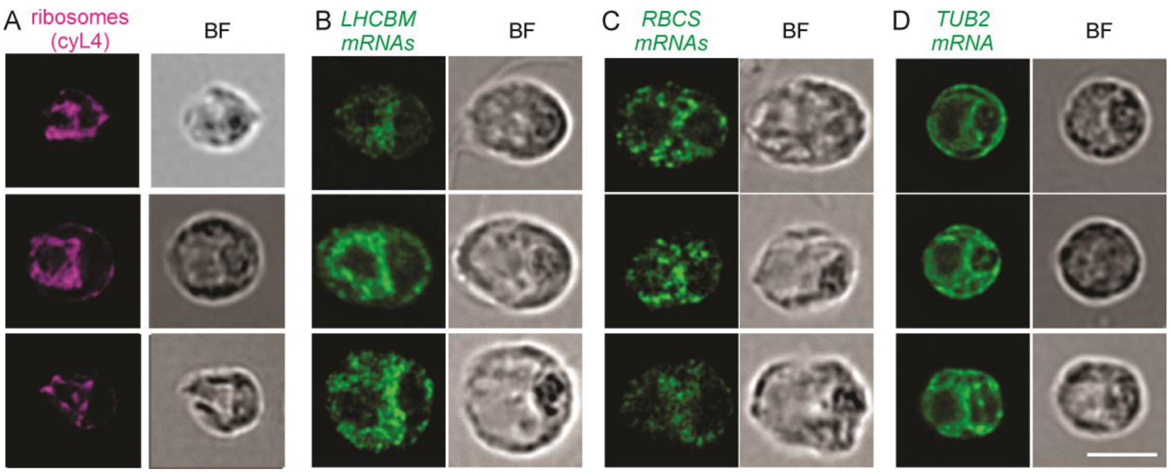

**Fig. S1.**
Localization patterns reported previously. Epifluorescence microscopy images of cells that were (A) IF-stained for cyL4 or (B-D) FISH-probed for (B) the LHCBM mRNAs, (C) the RBCS mRNAs or the (D) TUB2 mRNA. These cells show patterns that were reported previously (*7*, *8*); cyL4, LHCBM mRNAs and RBCS mRNAs, and TUB2 mRNA.  Error bar = 5.0 µm. Create a page break and paste in the Figure above the caption.

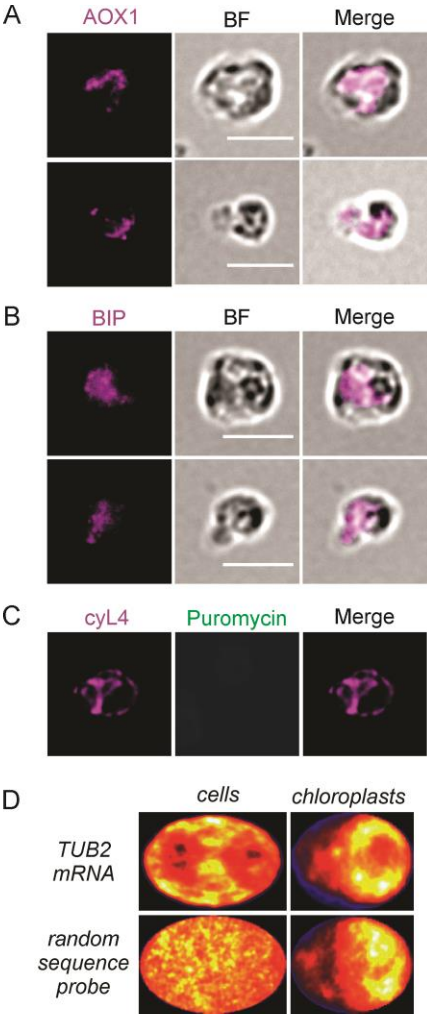

**Fig. S2.**
Experimental controls.  Isolated chloroplasts that were IF-stained for marker proteins for (A) mitochondria (AOX1) or (B) endoplasmic reticulum (BIP) do not show the localization pattern seen for (C) the ribosome marker protein (cyL4). (C) With the RPM method (Figure 3), the puromycin IF signal seen at the localized IF signal from cytoplasmic ribosomes (cyL4) on isolated chloroplasts is specific; it was not detected from chloroplasts that were not treated with puromycin. (D) The average distributions of the TUB2 mRNA FISH signal from all imaged cells and chloroplasts is compared to average distributions of the background signal from a control FISH probe with a random sequence that is not in the *Chlamydomonas* genome. The distributions of the average TUB2 mRNA and control FISH signals differ in cells, supporting the former as representing this mRNA. From chloroplasts, the signals are both weak and their distributions are similar, suggesting that much or all TUB2 mRNA FISH signal from chloroplasts is background

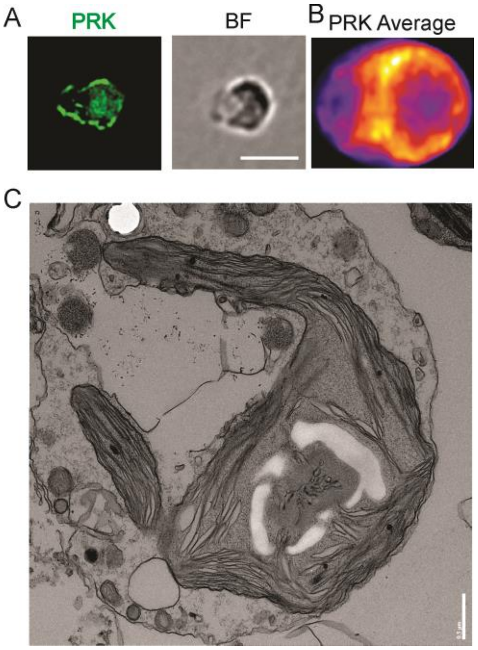

**Fig S3**

The isolated chloroplasts retain normal morphology (see Figure 1A for reference). (A) An isolated chloroplast IF stained for phosphoribulose kinase (PRK) to reveal the entire chloroplast and its retention of normal morphology during isolation. (B) A heatmap of the average PRK signal across all chloroplasts of the data set shows entire chloroplast and contrasts the heatmaps of the IF-signals of cyL4 (Figure 1D), puromycin (Figure 3C), and the FISH signals from the LHCB and RBCS mRNAs (Figure 4E) (n=30). (C) Transmission EM image of an isolated chloroplast shows that it has normal morphology. Cells were collected from cultures entrained to the diurnal cycle and processed as described previously (*15*). Images were acquired on a FEI Tecnai 12 120kv Transmission electron microscope using the Tecnai User Interface software and an AMTv601 CCD Camera. Settings used were an aperture of 3, a spot size of 2, and variable magnifications ranging from 2,900X to 68,000X.

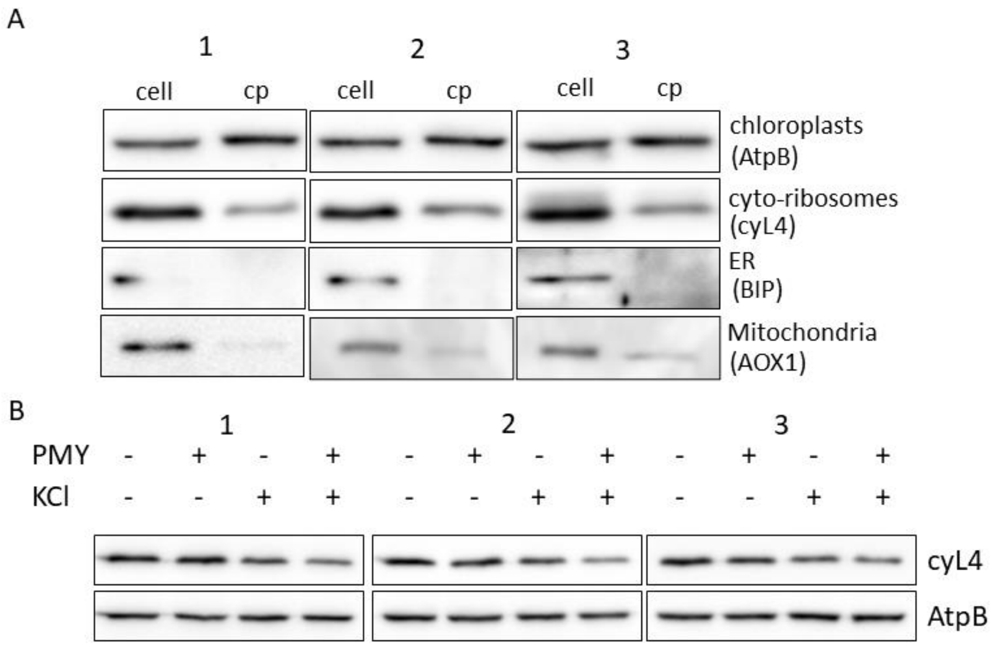

**Fig. S4.**
(A) These immunoblot results are represented by the bar heights in Figure 1B. Trials 1-3 are three biological replicate experiments, each performed from an independent culture. (B) These immunoblot results are represented by the bar heights in Figure 3C. Note the different order of treatments here and in the bar graph in Fig. 3D. Trials 1-3 are three biological replicate experiments, each performed from an independent culture and all conducted in parallel, including the immunoblots transfer, immune-reaction steps, and ECL imaging. KCl, incubation in 750 mM KCl; puromycin, PMY. The chloroplast protein AtpB was used as a loading control.

**Table S1. FISH probe sequences.**

| | | Probe Name | Probe Sequence (5’ to 3’) |
|---|---|---|---|
| oligo for labeling mRNA-specific probes | | FLAP X-Cy3 | Cy3/C ACT GAG TCC AGC TCG AAA CTT AGG AGG/Cy3<br><br>mRNA-specific probes below have a 3’ extension with the FLAPX reverse complement sequence:<br>CCTCCTAAGTTTCGAGCTGGACTCA<br>GTG |
| LHCBM | LHCBM7<br>Cre12.g548950<br><br>*≥95% sequence identity to the mRNA of LHCBM2<br>Cre12.g548400 | | |
| | | LHCII- 1 | GAGGACTTCATGATGGCGGCCATTTTGATTG-FLAPX reverse complement (above). |
| | | LHCII- 2* | AAGAAGCCGAACATGGAGAACATAGCCAGG C-FLAPX reverse complement (above). |
| | | LHCII- 3 | GTACATGCAGCTGCCGAGGGCCAAAAATTTA -FLAPX reverse complement (above). |
| | | LHCII- 4 | GCTCCTAAGCCTGTGAAAAGAGGCTCACACT-FLAPX reverse complement (above). |
| | | LHCII- 5* | GTCGCCCTCCGAGAAGGGGCCCAGGAACTTC -FLAPX reverse complement (above). |
| | | LHCII- 6* | GACAGACCGGCGGTGTCCCAGCCGTAGTCGC -FLAPX reverse complement (above). |
| | | LHCII- 7 | GATCAGCTCCAGCTCGCGGTAGCGCTTGAAG-FLAPX reverse complement (above). |
| | | LHCII- 8* | CCTTGAACCAGACAGCCTCACCGAACGGGAT -FLAPX reverse complement (above). |
| | | LHCII- 9* | AGGTAGTTCAGGCCGCCCTCAGCGAAGATCT-FLAPX reverse complement (above). |
| | | LHCII-10* | ATGATGGACTGGGCGTGGATCAGGTTCTCGT-FLAPX reverse complement (above). |
| | | LHCII-11* | TCAGCCAGGCCCATCACCACAACCTGGAAGG -FLAPX reverse complement (above). |
| | | LHCII-12* | ATCTCCTTCACCTTCAGCTCAGCGAAGGTGT-FLAPX reverse complement (above). |
| | | LHCII-13 | CGTAGCACCGCCACTTCGGTTAATCGCACGT-FLAPX reverse complement (above). |
| | | LHCII-14 | CAAAACCCGAACACAAAACTGAACCTCCGTA -FLAPX reverse complement (above). |
| | | LHCII-15 | GTGAACTTGGTGGCGTAGGCGAACGCGTTCA -FLAPX reverse complement (above). |
| | | LHCII-16* | TTGGCCAGGTGGTCGTCCAGGTTCTGGATGG-FLAPX reverse complement (above). |
| | | LHCII-17* | ATGCAGCCCAGAGCGCCCAGCATGGCCCAGC -FLAPX reverse complement (above). |
| | | LHCII-18 | ACCGGCTGCTCACGGTGGAGCGCACGGAGCT -FLAPX reverse complement (above). |

| | | | |
|---|---|---|---|
| | | LHCII-19 | CACCGCGCTCCTCTTCATCTCCGCTCAATCA-FLAPX reverse complement (above). |
| | | LHCII-20 | GTGGCCGTCAAGCCATTTTTAGTCTTCTCAA-FLAPX reverse complement (above). |
| | | LHCII-21 | TGCCGTGTTACACAACAAGGGCAAATCGCAA-FLAPX reverse complement (above). |
| | | LHCII-22 | TAGACAGCTAGAACAAAGCAGGCTGTAAAGA-FLAPX reverse complement (above). |
| | | LHCII-23 | TCAGCCAGGCCCAGGGGGTCAAAGGCACCAC-FLAPX reverse complement (above). |
| | | LHCII-24* | CCACTCGATGGCAGCGCGGGGCACCACGCGA-FLAPX reverse complement (above). |
| RBCS | RBCS2 Cre02.g120150 *≥95% sequence identity to the mRNA of RBCS1 (≥95% identity) Cre02.g120100 | | |
| | | RbcS- 1 | ACCCCATCAAACATCATCCTGGTTTGGCTGC-FLAPX reverse complement (above). |
| | | RbcS- 2 | GGCGGCCATTTTAAGATGTTGAGTGACTTCT-FLAPX reverse complement (above). |
| | | RbcS- 3* | GTCCAGACCATCATCTGGTTGGCCTGAGCCG-FLAPX reverse complement (above). |
| | | RbcS- 4* | GGTCCAGTAGCGGTTGTCGTAGTACAGCTGC-FLAPX reverse complement (above). |
| | | RbcS- 5* | ATGATCTGCACCTGCTTCTGGTTGTCGAAGG-FLAPX reverse complement (above). |
| | | RbcS- 6* | TTGGTGCAGGCGACGATCTCGCGCAGCACCT-FLAPX reverse complement (above). |
| | | RbcS- 7 | ACACGTAGGCCTTGTCCGACTCAGCGAACTC-FLAPX reverse complement (above). |
| | | RbcS- 8 | AATGTAGTCGACCTGGGCGGCGATCTGCTCG-FLAPX reverse complement (above). |
| | | RbcS- 9* | GCCACGGCCGCGGAGACGGAGGACTTGGCAA-FLAPX reverse complement (above). |
| | | RbcS-10 | TTACACGGAGCGCTTGTTGGCGGGCTGCCAG-FLAPX reverse complement (above). |
| | | RbcS-11* | TCGCGGCAGCCGAACATGGGCAGCTTCCACA-FLAPX reverse complement (above). |
| | | RbcS-12* | CAAGACACGCTGCCGAAGCGGATGGCCGACT-FLAPX reverse complement (above). |
| | | RbcS-13* | GCCTTGACGGCGGGCTTCAGCGCGGCCATGG-FLAPX reverse complement (above). |
| | | RbcS-14 | TAGGAGAAGGTCTCGAACATCTTGTTGTTGA-FLAPX reverse complement (above). |
| | | RbcS-15 | AGCAGTATCTTCCATCCACCGCCGTTCGTCA-FLAPX reverse complement (above). |
| | | RbcS-16 | GCACGAAACGGGGAGCTAAGCTACCGCTTCA-FLAPX reverse complement (above). |
| | | RbcS-17 | TGCAAAACTCCTCCGCTTTTTACGTGTTGAA-FLAPX reverse complement (above). |
| | | RbcS-18 | GGGGCAAGGCTCAGATCAACGAGCGCCTCCA-FLAPX reverse complement (above). |

| | | | |
|---|---|---|---|
| TUB2 | Cre12.g549550 | | |
| | | Tubulin-1 | CGATCACAAGCTCGAGTGGCCTGTGTAGAAG -FLAPX reverse complement (above). |
| | | Tubulin-2 | AAACCATGACGGCAAAAACATTATCAAGCAT -FLAPX reverse complement (above). |
| | | Tubulin-3 | TACGAAGAGTTCTTGTTCTGCACGTTCAGCA-FLAPX reverse complement (above). |
| | | Tubulin-4 | GCCTCCACACCAAAGCGTCAAATGGCAATCA -FLAPX reverse complement (above). |
| | | Tubulin-5 | CAGCTGCTATGGCCTATCACACAAGAGCTAA-FLAPX reverse complement (above). |
| Non-specific sequence | | Scramble | CTGAGTTAAGGCTTTCCACGGACGAGTTAAT-FLAPX reverse complement (above). |
